# Immune-informed mucin hydrogels evade fibrotic foreign body response *in vivo*

**DOI:** 10.1101/554865

**Authors:** Hongji Yan, Cédric Seignez, Morgan Hjorth, Benjamin Winkeljann, Oliver Lieleg, Mia Phillipson, Thomas Crouzier

## Abstract

The immune-mediated foreign body response to biomaterial implants can trigger the formation of insulating fibrotic capsules that can compromise implant function. To address this challenge, we leverage the intrinsic bioactivity of the mucin biopolymer, a heavily glycosylated protein that forms the protective mucus gel covering mucosal epithelia. By using a bioorthogonal inverse electron demand Diels-Alder reaction, we crosslink mucins into implantable hydrogels. We show that mucin hydrogels (Muc-gels) modulate the immune response driving biomaterial-induced fibrosis. Muc-gels did not elicit fibrosis 21 days after implantation in the peritoneal cavity of C57Bl/6 mice, whereas medical-grade alginate hydrogels (Alg-gels) were covered by fibrous tissues. Further, Muc-gels dampened the recruitment of innate and adaptive immune cells to the gel and triggered a pattern of very mild activation marked by a noticeably low expression of the fibrosis-stimulating TGF-β1 cytokine. With this advance in mucin materials, we provide an essential tool to better understand mucin bioactivities and to initiate the development of new mucin-based and mucin-inspired ‘immune-informed’ materials for implantable devices subject to fibrotic encapsulation.

## Introduction

An universal challenge for functional implants such as encapsulated cells^1^, sensors^2^, electrodes^3^, and drug-eluting materials^4^ is their rapid isolation from the body by a fibrotic capsule mediated by a foreign body reaction (FBR). The FBR starts immediately after implantation with the passive absorption of proteins onto the implant. This is followed by accumulation of monocytes and macrophages, that can further fuse to form multinucleated giant cells. The subsequent secretion of fibroblast-recruiting and pro-angiogenic factors by activated immune cells leads to the generation of a fibrous capsule around the biomaterial 2 to 4 weeks after implantation^5, 6^. To prevent fibrosis, biomaterials can be designed to reduce the non-specific binding of proteins and subsequent cell adhesion, or release anti-inflammatory drugs such as glucocorticoid dexamethasone^7, 8^ that dampen the immune response to implants. However, these strategies are relatively short-lived. Alternative approaches, which consist in careful physical and molecular design of materials, appear as promising strategies to modulate the immune response to implants since it can provide a more durable and resilient immune modulating signal. For instance, physical properties of materials such as size^1^, geometry^9^, roughness^10^, porosity^11^, hydrophobicity^12^, mechanical properties^13^ and chemistry^14^ have all shown to influence immune activation.

Another strategy is to leverage mechanisms by which the immune system distinguishes foreign objects from the body’s own components. For instance, bacteria-derived LPS-presenting objects are marked as foreign and readily initiate immune responses, while cell surface markers such as CD47^15^ and CD200^16^ act as self-signalling molecules and contribute to lowering the immune activation state. Materials conjugated with CD47 could prevent macrophage attachment, activation, and phagocytosis^17, 18^, while polystyrene grafted with CD200 decreased the expression of TNF-α and IL 6 in macrophages^19^.

Mucin glycoproteins have emerged as another important part of the immune regulatory arsenal in mammals. Mucins are either secreted to form the mucus gel that covers the mucosal epithelia, or expressed as cell transmembrane proteins as a main component of the cell glycocalyx. Every protein of the mucin family features densely glycosylated serine and threonine-rich regions. The mostly O-linked glycans are terminated by a wide variety of sugars, with sialic acids and fucose residues being the most common. Owing to this rich chemistry, the mucus gel is a multifunctional material that hydrates, lubricates and forms selective barriers that protect the epithelium. Secreted mucins have shown an immunological role, although their exact activity is still unclear. For instance, in one study, intestinal MUC2 mucins glycans were shown to dampen inflammatory activation of dendritic cells as evidenced by the increased IL- 10 secretion^20^, while a more recent study suggested the same mucin can increase the expression of proinflammatory cytokine IL-8 in dendritic cells^21^. Cancerous cells can escape the immune attack by expressing membrane-bound MUC1 mucins with altered glycosylation and by secreting substantial amounts of mucins that form a physical and biochemical protective encasing of the tumor^22^. Mucins-containing materials have been suggested as potential building blocks for multifunctional biomaterials, but their immune modulating properties have not been investigated^23, 24^.

Given the strong indications that mucins hold immune-modulatory capacities, we aimed to develop a mucin material that can deliver immunological signals upon implantation. We show that “clickable” bovine submaxillary mucin (BSM) can be assembled into a robust hydrogel material (Muc-gel). Muc-gels implanted into the peritoneal cavity of mice showed no fibrosis after 21 days, while alginate hydrogels (Alg-gel) were covered by a fibrotic capsule. Analysis of the immune activity on and around the implant revealed large differences in cell composition and activation when compared to an Alg-gel, suggesting strong immune-modulating properties of this new class of material.

### Click mucins can be assembled into implantable hydrogels

We successfully introduced either tetrazine (Tz) or norbornene (Nb) ‘clickable’ functionality to BSM molecules by reacting Tz and Nb amine derivatives with activated carboxylic groups of mucins (**Figure 1A**) as confirmed by^1^ H NMR (**Figure S-1**). The resulting BSM-Tz and BSM-Nb form covalent bonds through an inverse electron demand Diels-Alder cycloaddition reaction when mixed in solution (**Figure 1B**). This bio-orthogonal reaction has no known reactivity with natural molecules and is thus suitable for cell encapsulation and *in vivo* implantation^25^. Time-sweeping rheology analysis revealed that the sample transitioned from a viscous solution (*G*’’ > *G*’) to a gel dominated by the elastic modulus after ~ 4 min then increased dramatically within 20 min and slowly reached a plateau-like state after ~ 40 min (**Figure 1C**). A frequency sweep performed after the viscoelastic moduli have reached plateaus demonstrates that the system appears indeed to be efficiently and covalently cross-linked (**Figure 1C**). Based on the rheological data and rubber elasticity theory^26, 27^, we estimate the distance between two adjacent crosslinkers (𝝃) to be ~ 7,4 nm (see methods in SI), which would allow for unrestricted diffusion of nutrients and cell secretomes through the gels^28^. Indeed, THP-1 derived macrophage type 0 embedded in the Muc-gel maintained their viability over a 7-day culture (**Figure S-2**). Muc-gels were degraded by proteases as indicated by a gradual decrease of the Muc-gels’ storage modulus when exposed to a trypsin solution (**Figure 1D**). This indicates that the Muc-gel is primarily linked through the non-glycosylated protein backbone, which is sensitive to trypsin (**Figure 1E**). This is also supported by the unchanged sialic acid content measured before and after functionalisation of the mucins (**Figure S-3**). Such a versatile system allows for the rapid assembly of mucin molecules into robust hydrogels. Those gels are useful as model materials to study mucin bioactivity, as the local immune-isolation for cell transplantation, and as implantable materials or implant coatings.

**Figure 1.**
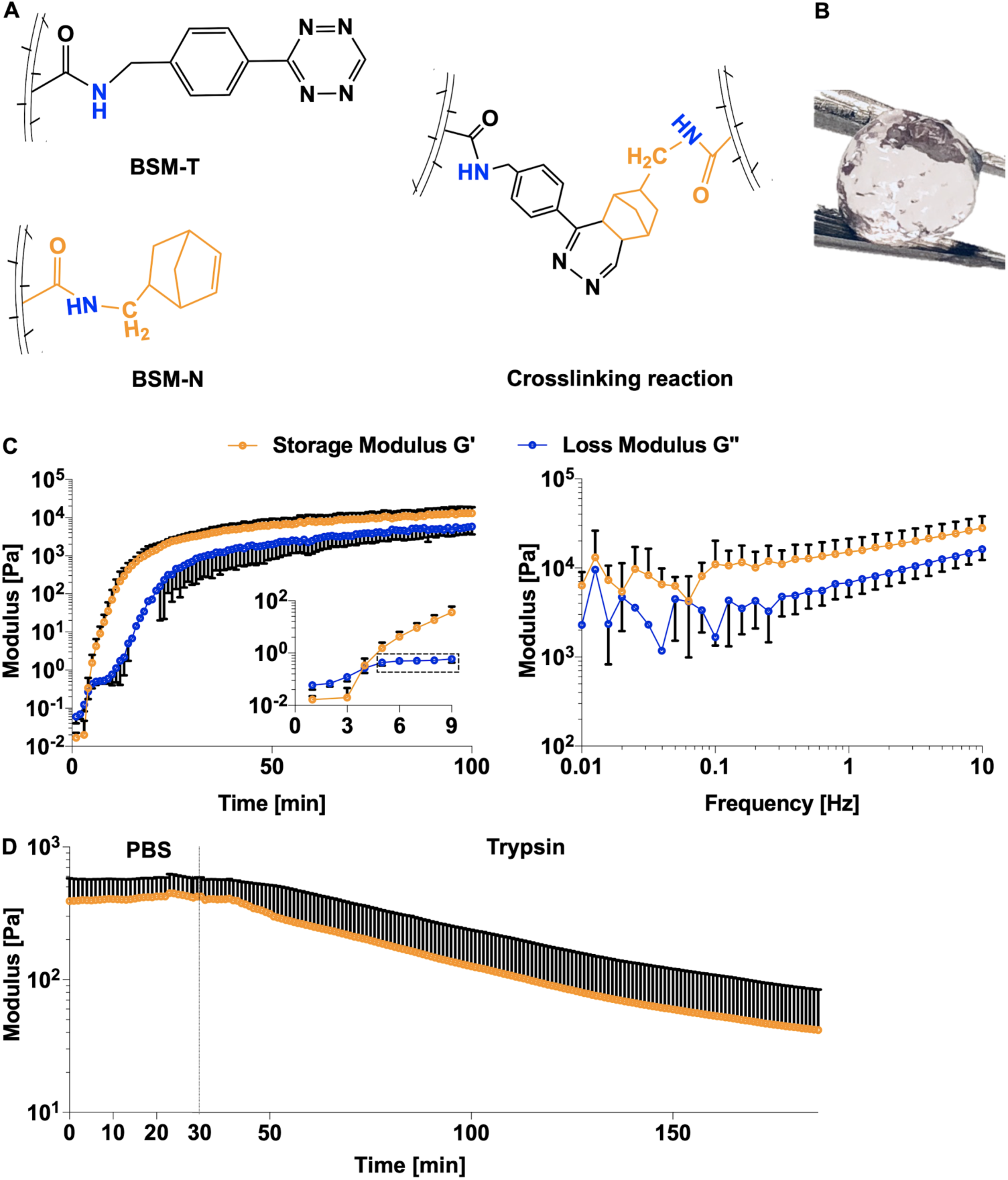
Synthesis and rheological properties of Muc-gels. (A) Tetrazine (Tz) and norbornene (Nb) conjugated BSM and the crosslinking reaction. (B) Mixing solutions of BSM-Tz and BSM-Nb leads to the formation of a robust Muc-gel. (C) Left: Time-dependent rheological measurements of the mixed BSM-Tz and BSM-Nb solution demonstrate the formation of a Muc-gel after ~4 minutes (insert). Right: Final frequency dependent viscoelastic moduli of the cross-linked Muc-gel. (D) Trypsin-induced Muc-gel degradation. After gelation, the Muc-gel was first exposed to PBS and then to a trypsin solution. In all graphs, the error bars denote the standard deviation as obtained from measurements of *n* = 3 independent samples.

### Muc-gels implanted into the peritoneal cavity of mice averted fibrotic encapsulation

The foreign body reaction (FBR) to Muc-gels was investigated by using immunocompetent C57Bl/6 mice in which the FBR is in many respects similar to that observed in humans^1, 29^. We implanted customized Alg-gel cylinders (diameter=4.78 mm, height= 2,79 mm, 50 µl, **Figure 1A**) or Muc-gel cylinders (**Figure 1B**) into the peritoneal cavity (i.p.) and compared the outcome on day 21, as Alg-gel cylinders are known to trigger fibrosis when implanted into C57Bl/6 mice^29^. After 21 days of implantation, we found Alg-gels were surrounded by clear fibrous capsules (**Figure 2A, a-b)**, whereas there was no sign of fibrosis on the surface of Muc-gels, as illustrated by the few cells at the gel surface stained by H&E (**Figure 2B, c-d**). Masson’s trichrome staining indicated the presence of a collagen-rich matrix in the tissue formed around Alg-gels (**Figure 2e**, **f**, blue arrows), whereas the Muc-gels showed no staining (**Figure 2g**, **h**). These results demonstrated that Muc-gels prevent the development of fibrosis *in vivo*.

**Figure 2.**
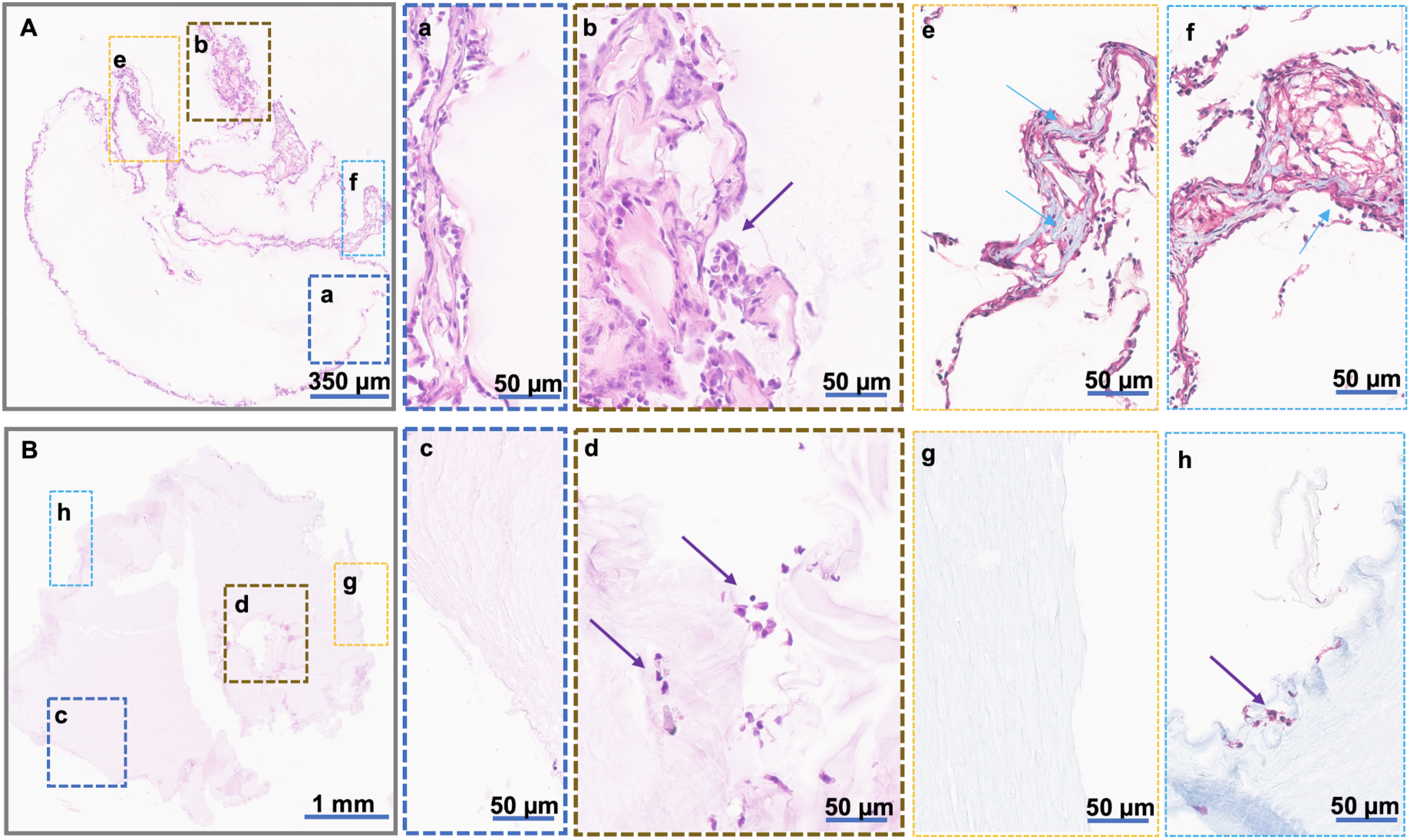
Comparing the fibrotic capsule formation on Alg-gels (A) and Muc-gels (B) that were harvested 21 days post-i.p. implantation in mice by histological evaluation. (A, B) Hematoxylin and Eosin (H&E) staining. (a-b, c-d) H&E staining of magnified region indicated in A and B. (e-f, g-h) Masson’s trichrome staining of magnified regions indicated in A and B. Arrows indicate cells (purple) or collagen revealed by Masson’s trichrome staining (blue). Data are from 4 mice (n=12).

### Muc-gel implants recruited fewer macrophages and B cells

To investigate the cellular mechanism by which Muc-gels averted fibrotic capsule formation, we characterized the local immune response to the implanted materials. We examined both innate and adaptive immune cell compositions in the peritoneal cavity by performing lavages, and from Muc-gels and Alg-gels implants retrieved on day 14 and 21 (**Figure 3**). Interestingly, the cell populations in the peritoneal cavity reflected rather closely the ones found associated with explanted hydrogels. Both on day 14 and 21, myeloid cells (CD68^−^Gr1^low/-^CD11b^+^) represented a majority of the cell population detected by FACS, followed by macrophages (CD68^+^CD11b^+^), B lymphocytes (CD19^+^IgM^+^) and neutrophils (CD11b^+^Gr1^hi^). Neutrophils, which can typically be found hours to weeks after implantation^30–32^, were here found as a very low proportion at these later time points. The proportion of macrophages and B cells was significantly lower in Muc-gels compared to Alg-gels on day 14. On day 21, when the complete fibrotic isolation occurred, the proportion of macrophages and B cells dropped in mice with implanted Alg-gels to levels comparable to those of mice with Muc-gels. B cells represented a small but significant population of the immune cells present in the i.p. space at 2 weeks following implantation of Alg-gels but not Muc-gels. Although their contribution to the FBR is unclear, it has been recently shown that B cells could potentiate the biomaterial-induced fibrosis, perhaps by regulating macrophage phenotype and cytokine secretion^33, 34^.

**Figure 3.**
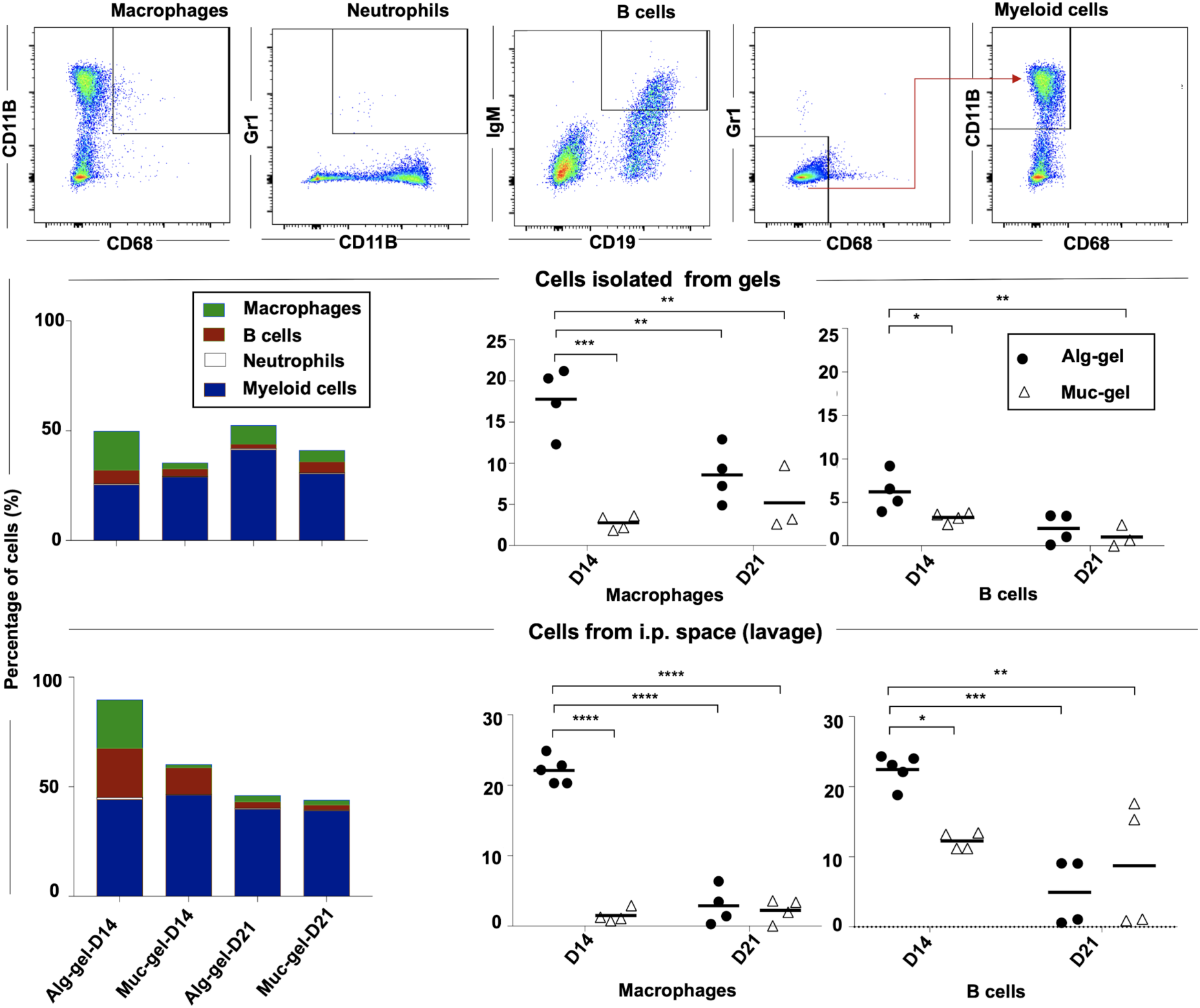
The host immune response to implanted Alg-gels and Muc-gels *in vivo*. A representative FACS profile showing the gating strategy used for the analysis of host macrophages (CD68^+^CD11b^+^), neutrophils (CD11b^+^Gr1^+^), B cells (CD19^+^IgM^+^), and myeloid cells (CD68^−^Gr1^low/-^CD11b^+^). Cell frequency of host immune cells to mononuclear cells retrieved from the i.p. space (lavage), or from gels on day 14 and 21 post-implantation in the mice. Data are from 4 or 5 mice. Differences were determined using One-way ANOVA with Bonferroni-correction.

The presence of immune cells on Muc-gels suggests that the antifibrotic effect is not only achieved by preventing cell to recognize and interact with the Muc-gels. Rather the Muc-gel dampened the immune cell recruitment.

### Muc-gel implants dampened the macrophage and myeloid cell activation

Chemokines and cytokines produced by activated immune cells play essential roles in the FBR cascades, i.e., inducing cell recruitment and differentiation of other immune cells and fibroblasts into myofibroblasts that participate in fibrotic capsule formation. We focused our attention on macrophages (CD68^+^CD11b^+^) and other myeloid cells (CD68^−^Gr1^low/-^CD11b^+^) that were predominant fractions of immune cells sorted by FACS (**Figure 4, S-4**). Macrophages are especially relevant in this context since they display remarkable functional plasticity and participate in the host defence, immune regulation, and tissue repair, all aspects playing a pivotal role in mediating implant-triggered fibrosis^3, 34, 35^.

**Figure 4.**
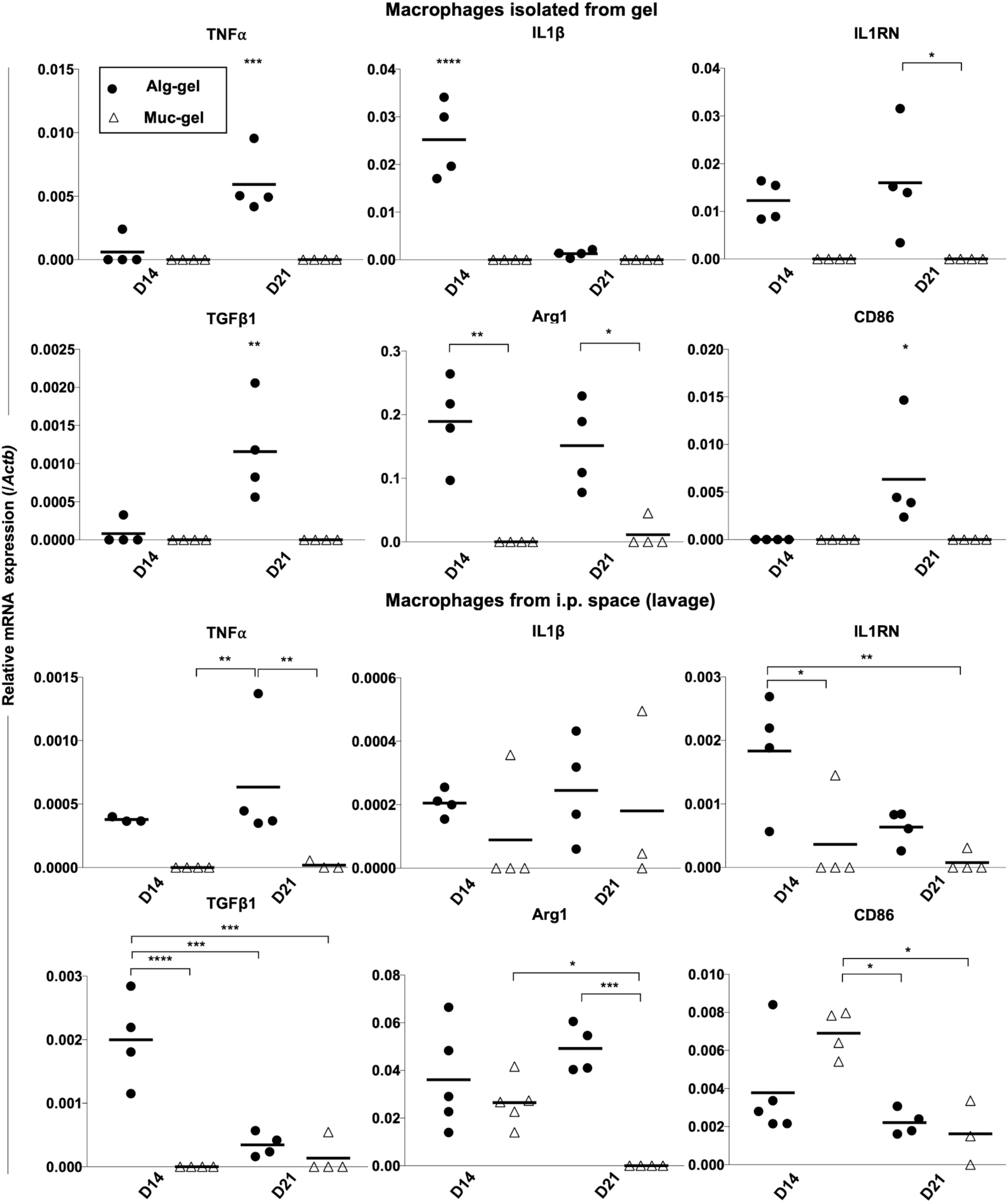
Muc-gels dampened the macrophage activation. Gene expression in freshly sorted macrophages (CD68^+^CD11b^+^) from the explant and i.p. space on day 14 and 21 was analysed by RT-PCR. Data represents the mean of duplicates of RT-PCR from 3 to 5 independent experiments and from four mice. Differences were determined using a One-way ANOVA with Bonferroni-correction.

Strikingly, at 14 and 21 days’ post-implantation, the macrophages from Muc-gels and the corresponding peritoneal cavity showed nearly no expression of the analysed pro-inflammatory and pro-fibrotic markers^36, 37, 38^ (**Figure 4)**. This was in strong contrast with the macrophages from Alg-gels, which expressed significant higher levels of M1 marker (pro-inflammatory cytokines and cell surface molecule) TNF-α and CD86 on day 21 and IL-1β on day 14, and of the M2 markers (anti-inflammatory cytokines and intracellular enzyme) IL-1RN on day 21, TGF-β1 on day 14, and Arg1 on day 14 and 21. The cytokine and cell-marker expression signature of the myeloid cells (CD68^−^Gr1^low/-^CD11b^+^) were in many aspects similar to the pattern found for macrophages (**Figure S-4)**. Exceptions were for TNF-α and CD86 that were found to be expressed at significantly higher levels on day 14 in cells from the Muc-gels and their i.p. space compared to Alg-gels.

Although chemokines and cell-surface markers can, in principle, discriminate between macrophages of inflammatory (M1) or alternatively activated (M2) phenotype, no distinct signature could be found in our study. This is exemplified by the simultaneous expression of typical M1 and M2 markers in macrophages on and around Alg-gels. It is now understood that rather than being at distinct activation states, macrophages *in vivo* are found on a continuum between the classic M1 versus alternative M2 phenotypes^39^. Our results reinforce the observation that biomaterials implanted in the peritoneal cavity of mice lead to macrophages with both M1 and M2 characteristics^40, 41^. Importantly and regardless of the nomenclature, several cytokines known for their roles in non-biomaterial-induced fibrosis were expressed by immune cells on and around Alg-gels. For instance, TNF-α is a critical factor in the liver and pulmonary fibrosis^42^, IL-1β was also found to potentiate the pulmonary fibrosis^43, 44^, and the ratio of IL-1β to IL-1RN expression determines its extent^45^. In addition, Arg1, an enzyme expressed in M2 macrophages promotes collagen deposition, which has also been associated with fibrosis^46^. Finally, TGF-β1 is strongly linked to tissue fibrotic cascades^47, 48^ and biomaterial-induced fibrosis^49, 50^. At high concentrations, TGF-β1 can increase the extracellular matrix deposition by stimulating the differentiation of quiescent fibroblasts to collagen type-I producing myofibroblasts. Remarkably, none of these fibrosis-related cytokines and chemokines were expressed by macrophages and myeloid cells in contact with Muc-gels. It could thus be that the Muc-gels provide the necessary signals for macrophages and other immune cells to adopt a ‘fibrosis-repressing’ phenotype.

### Muc-gel implants showed no TGF-β1 and α-smooth muscle actin protein expression

Next, we studied protein expression of TGF-β1 and α-smooth muscle actin, two markers of biomaterial-induced fibrosis^50^, by immunohistological staining of day-21 explants (**Figure 5)**. The Muc-gel surface had very few cells as indicated by DAPI staining, whereas Alg-gels were surrounded by a few layers of cells, which is in agreement with the H&E staining (**Figure 2A**, **B**). The strong expression of α-smooth muscle actin in the tissue surrounding of Alg-gels confirmed its fibrotic nature (**Figure 5 red**). In addition, TGF-β1 expression was induced by Alg-gels (**Figure 5 green**), whereas it was not expressed around and on the Muc-gel. This is in agreement with the low TGF-β1 gene expression levels measured in macrophages and myeloid cells (CD68^−^Gr1^low/-^CD11b^+^) (**Figures 4 and S-4**). The absence of TGF-β1 is likely to contribute to the absence of fibrotic capsule around Muc-gels.

**Figure 5.**
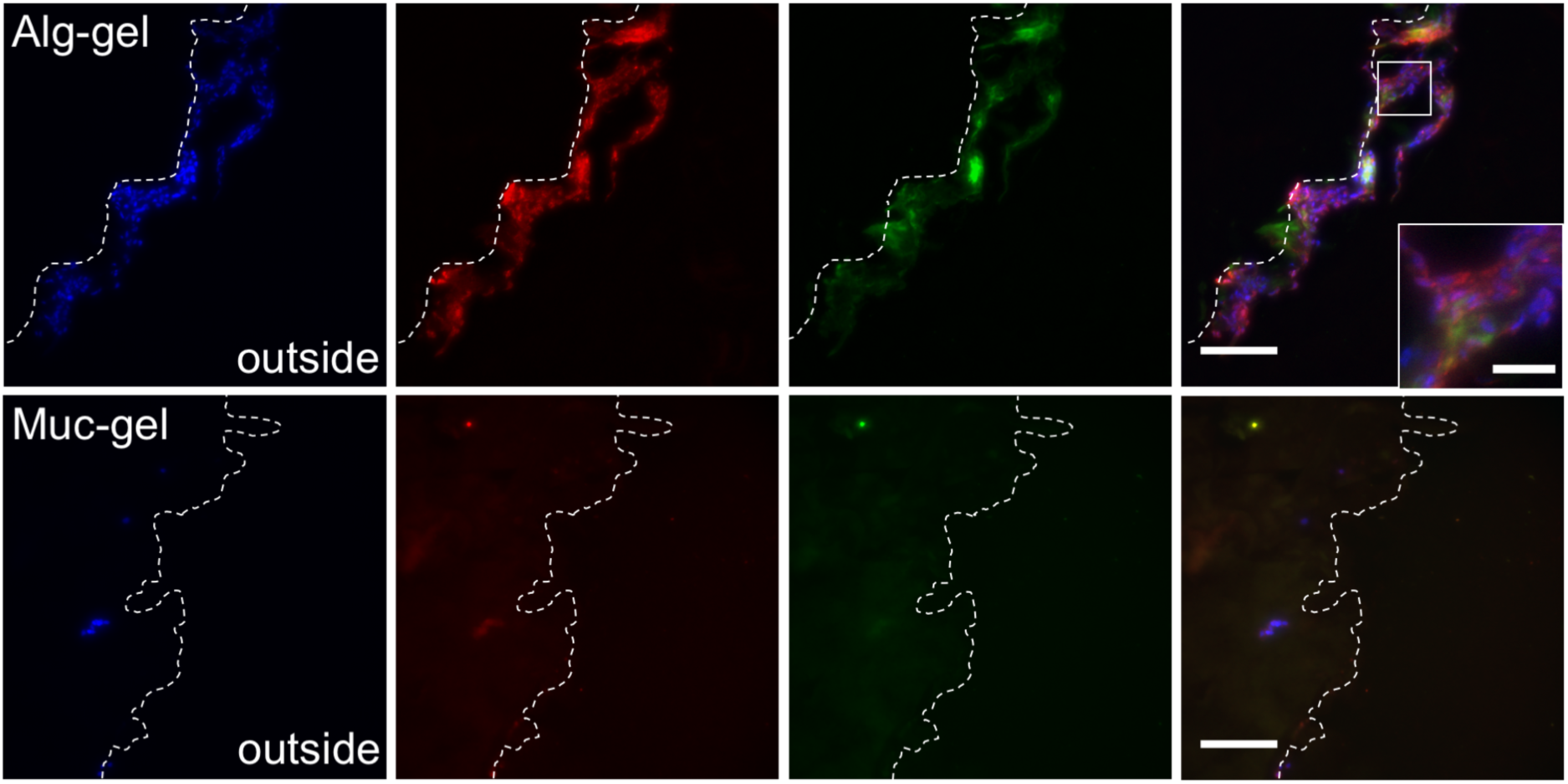
Immunohistofluorescence staining of explanted day-21 Alg-gels and Muc-gels. In blue nuclei (DAPI), in red α-smooth muscle actin, a marker for fibrosis, and in green TGF- β1. The zoomed-in insert for Alg-gel shows the localization of α-smooth muscle actin in the periphery of the nuclei and the presence of TGF-β1 in the cytoplasm and extracellular space. The dotted white line indicates the interface of the hydrogels. Scale bars are 100 µm, and 25 µm for the insert. n=4 from four mice.

We chose BSM as the model mucin for this study because it is well-characterized with known amino acids^51^ and carbohydrate components. The main O-linked glycan on the BSM is the sialyl Tn antigen^52^, which is composed of N-acetylgalactosamine linked to a terminal sialic acid residue via a α2-6 glycosidic bond. The sialic acid residue of BSM binds to several sialic acid-binding immunoglobulin superfamily lectins (siglec) 2, 3, 5, 6, and 9^53, 54^, which regulate the function of macrophages, B cells, neutrophils, monocytes and others immune cells^55^ and could explain some of the effects observed here. Sialylated glycans are recognized as ‘self’ ligands by siglec and tend to dampen an excessive innate immune response^56^. However, protein fragments on the mucin backbone could also be important to elicit such bioactivities.

## Conclusion

We have developed a first ‘clickable’ mucin material that is capable of modulating the immune response to avert biomaterial-induced fibrosis. Our results highlight the strong bioactivity of mucin molecules and their potential value as building blocks of immune-modulatory materials. The mucin hydrogel is biocompatible and can be assembled into various forms (macro-gel, droplets, thin films) opening a variety of applications including cell transplantations and biomaterial surface modifications. This work also contributes to shifting the perception of extracellular mucins and mucus away from a passive barrier material sitting on our mucosal epithelium towards a matrix rich in biochemical signals.

## Acknowledgements

OL acknowledges financial support from the Deutsche Forschungsgemeinschaft (DFG) through project B11 in the framework of SFB863. TC acknowledges financial support from the Swedish Foundation for Strategic Research (grant FFL15-0072), FORMAS (grant 2015- 1316), and the Swedish Research Council (grant 2014-6203). MP is supported by the Swedish Research Council, Ragnar Söderberg foundation and Knut and Alice Wallenberg foundation. The authors acknowledge Xueying Zhong for her participation in preliminary work on the ‘clickable’ mucin hydrogel, and the histology resources from Department of Pathology and Wildlife Diseases, National Veterinary Institute (SVA) and from Scilifelab Tissue Profiling Facility at Uppsala University.

## References

1. Veiseh, O. et al. Size- and shape-dependent foreign body immune response to materials implanted in rodents and non-human primates. Nat. Mater. 14, 643–651 (2015).

2. Avula, M. N., Rao, A. N., McGill, L. D., Grainger, D. W. & Solzbacher, F. Modulation of the foreign body response to implanted sensor models through device-based delivery of the tyrosine kinase inhibitor, masitinib. Biomaterials 34, 9737–9746 (2013).

3. Lotti, F., Ranieri, F., Vadalà, G., Zollo, L. & Di Pino, G. Invasive Intraneural Interfaces: Foreign Body Reaction Issues. Front. Neurosci. 11, 497 (2017).

4. Kearney, C. J. & Mooney, D. J. Macroscale delivery systems for molecular and cellular payloads. Nat. Mater. 12, 1004–1017 (2013).

5. Sridharan, R., Cameron, A. R., Kelly, D. J., Kearney, C. J. & O’Brien, F. J. Biomaterial based modulation of macrophage polarization: a review and suggested design principles. Mater. Today 18, 313–325 (2015).

6. Anderson, J. M., Rodriguez, A. & Chang, D. T. Foreign body reaction to biomaterials. Semin. Immunol. 20, 86–100 (2007).

7. Hickey, T., Kreutzer, D., Burgess, D. J. & Moussy, F. Dexamethasone/PLGA microspheres for continuous delivery of an anti-inflammatory drug for implantable medical devices. Biomaterials 23, 1649–1656 (2002).

8. Daugherty, A. L., Cleland, J. L., Duenas, E. M. & Mrsny, R. J. Pharmacological modulation of the tissue response to implanted polylactic-co-glycolic acid microspheres. Eur. J. Pharm. Biopharm. 44, 89–102 (1997).

9. Matlaga, B. F., Yasenchak, L. P. & Salthouse, T. N. Tissue response to implanted polymers: the significance of sample shape. J. Biomed. Mater. Res. 10, 391–397 (1976).

10. Wójciak-Stothard, B., Curtis, A., Monaghan, W., MacDonald, K. & Wilkinson, C. Guidance and activation of murine macrophages by nanometric scale topography. Exp. Cell Res. 223, 426–435 (1996).

11. Sussman, E. M., Halpin, M. C., Muster, J., Moon, R. T. & Ratner, B. D. Porous implants modulate healing and induce shifts in local macrophage polarization in the foreign body reaction. Ann. Biomed. Eng. 42, 1508–1516 (2014).

12. Bhaskar, V., Bhuvanesh, T., Ma, N., Lendlein, A. & Roch, T. The interaction of human macrophage subsets with silicone as a biomaterial. Clin. Hemorheol. Microcirc. 61, 119–133 (2015).

13. Patel, N. R. et al. Cell elasticity determines macrophage function. PLoS One 7, e41024 (2012).

14. Barbosa, J. N., Madureira, P., Barbosa, M. A. & Aguas, A. P. The influence of functional groups of self-assembled monolayers on fibrous capsule formation and cell recruitment. Journal of Biomedical Materials Research Part A: An Official Journal of The Society for Biomaterials, The Japanese Society for Biomaterials, and The Australian Society for Biomaterials and the Korean Society for Biomaterials 76, 737–743 (2006).

15. Jiang, P., Lagenaur, C. F. & Narayanan, V. Integrin-associated protein is a ligand for the P84 neural adhesion molecule. J. Biol. Chem. 274, 559–562 (1999).

16. Hoek, R. M. et al. Down-regulation of the macrophage lineage through interaction with OX2 (CD200). Science 290, 1768–1771 (2000).

17. Tsai, R. K. & Discher, D. E. Inhibition of ‘self’ engulfment through deactivation of myosin-II at the phagocytic synapse between human cells. J. Cell Biol. 180, 989–1003 (2008).

18. Stachelek, S. J. et al. The effect of CD47 modified polymer surfaces on inflammatory cell attachment and activation. Biomaterials 32, 4317–4326 (2011).

19. Kim, Y. K., Que, R., Wang, S.-W. & Liu, W. F. Modification of biomaterials with a self-protein inhibits the macrophage response. Adv. Healthc. Mater. 3, 989–994 (2014).

20. Shan, M. et al. Mucus enhances gut homeostasis and oral tolerance by delivering immunoregulatory signals. Science 342, 447–453 (2013).

21. Melo-Gonzalez, F. et al. Intestinal mucin activates human dendritic cells and IL-8 production in a glycan-specific manner. J. Biol. Chem. (2018). doi:10.1074/jbc.M117.789305

22. Hollingsworth, M. A. & Swanson, B. J. Mucins in cancer: protection and control of the cell surface. Nat. Rev. Cancer 4, 45–60 (2004).

23. Petrou, G. & Crouzier, T. Mucins as multifunctional building blocks of biomaterials. Biomaterials Science (2018). doi:10.1039/c8bm00471d

24. Sandberg, T., Carlsson, J. & Ott, M. K. Mucin coatings suppress neutrophil adhesion to a polymeric model biomaterial. Microsc. Res. Tech. 70, 864–868 (2007).

25. Madl, C. M. & Heilshorn, S. C. Bioorthogonal Strategies for Engineering Extracellular Matrices. Adv. Funct. Mater. 28, 1706046 (2018).

26. Yan, H. J. et al. Synthetic design of growth factor sequestering extracellular matrix mimetic hydrogel for promoting in vivo bone formation. Biomaterials 161, 190–202 (2018).

27. Welzel, P. B. et al. Modulating Biofunctional starPEG Heparin Hydrogels by Varying Size and Ratio of the Constituents. Polymers 3, 602–620 (2011).

28. Cruise, G. M., Scharp, D. S. & Hubbell, J. A. Characterization of permeability and network structure of interfacially photopolymerized poly(ethylene glycol) diacrylate hydrogels. Biomaterials 19, 1287–1294 (1998).

29. Kolb, M. et al. Differences in the fibrogenic response after transfer of active transforming growth factor-beta1 gene to lungs of ‘fibrosis-prone’ and ‘fibrosis-resistant’ mouse strains. Am. J. Respir. Cell Mol. Biol. 27, 141–150 (2002).

30. Anderson, J. M. Biological Responses to Materials. Annu. Rev. Mater. Res. 31, 81–110 (2001).

31. Mantovani, A., Cassatella, M. A., Costantini, C. & Jaillon, S. Neutrophils in the activation and regulation of innate and adaptive immunity. Nat. Rev. Immunol. 11, 519–531 (2011).

32. Jhunjhunwala, S. et al. Neutrophil Responses to Sterile Implant Materials. PLoS One 10, e0137550 (2015).

33. Affara, N. I. et al. B cells regulate macrophage phenotype and response to chemotherapy in squamous carcinomas. Cancer Cell 25, 809–821 (2014).

34. Doloff, J. C. et al. Colony stimulating factor-1 receptor is a central component of the foreign body response to biomaterial implants in rodents and non-human primates. Nat. Mater. 16, 671–680 (2017).

35. Dondossola, E. et al. Examination of the foreign body response to biomaterials by nonlinear intravital microscopy. Nat Biomed Eng 1, (2016).

36. Gordon, S. & Martinez, F. O. Alternative activation of macrophages: mechanism and functions. Immunity 32, 593–604 (2010).

37. Mosser, D. M. & Edwards, J. P. Exploring the full spectrum of macrophage activation. Nat. Rev. Immunol. 8, 958–969 (2008).

38. Martinez, F. O., Sica, A., Mantovani, A. & Locati, M. Macrophage activation and polarization. Front. Biosci. 13, 453–461 (2008).

39. Mantovani, A., Sozzani, S., Locati, M., Allavena, P. & Sica, A. Macrophage polarization: tumorassociated macrophages as a paradigm for polarized M2 mononuclear phagocytes. Trends Immunol. 23, 549–555 (2002).

40. Moore, L. B., Sawyer, A. J., Charokopos, A., Skokos, E. A. & Kyriakides, T. R. Loss of monocyte chemoattractant protein-1 alters macrophage polarization and reduces NFκB activation in the foreign body response. Acta Biomater. 11, 37–47 (2015).

41. Mooney, J. E. et al. Transcriptional switching in macrophages associated with the peritoneal foreign body response. Immunol. Cell Biol. 92, 518–526 (2014).

42. Piguet, P. F., Ribaux, C., Karpuz, V., Grau, G. E. & Kapanci, Y. Expression and localization of tumor necrosis factor-alpha and its mRNA in idiopathic pulmonary fibrosis. Am. J. Pathol. 143, 651–655 (1993).

43. Gasse, P. et al. IL-1R1/MyD88 signaling and the inflammasome are essential in pulmonary inflammation and fibrosis in mice. J. Clin. Invest. 117, 3786–3799 (2007).

44. Kolb, M., Margetts, P. J., Anthony, D. C., Pitossi, F. & Gauldie, J. Transient expression of IL-1β induces acute lung injury and chronic repair leading to pulmonary fibrosis. J. Clin. Invest. 107, 1529–1536 (2001).

45. Arend, W. P. Interleukin-1 Receptor Antagonist. in Advances in Immunology (ed. Dixon, F. J.) 54, 167–227 (Academic Press, 1993).

46. Barron, L. & Wynn, T. A. Macrophage activation governs schistosomiasis-induced inflammation and fibrosis. Eur. J. Immunol. 41, 2509–2514 (2011).

47. Border, W. A. & Noble, N. A. Transforming Growth Factor β in Tissue Fibrosis. N. Engl. J. Med. 331, 1286–1292 (1994).

48. Wynn, T. A. Fibrotic disease and the T(H)1/T(H)2 paradigm. Nat. Rev. Immunol. 4, 583–594 (2004).

49. Kenneth Ward, W. A review of the foreign-body response to subcutaneously-implanted devices: the role of macrophages and cytokines in biofouling and fibrosis. J. Diabetes Sci. Technol. 2, 768–777 (2008).

50. Li, A. G. et al. Elevation of transforming growth factor beta (TGFbeta) and its downstream mediators in subcutaneous foreign body capsule tissue. J. Biomed. Mater. Res. A 82, 498–508 (2007).

51. Jiang, W., Gupta, D., Gallagher, D., Davis, S. & Bhavanandan, V. P. The central domain of bovine submaxillary mucin consists of over 50 tandem repeats of 329 amino acids. Eur. J. Biochem. 267, 2208–2217 (2000).

52. Tsuji, T. & Osawa, T. Carbohydrate structures of bovine submaxillary mucin. Carbohydr. Res. 151, 391–402 (1986).

53. Macauley, M. S., Crocker, P. R. & Paulson, J. C. Siglec-mediated regulation of immune cell function in disease. Nat. Rev. Immunol. 14, 653–666 (2014).

54. Ohta, M. et al. Immunomodulation of monocyte-derived dendritic cells through ligation of tumor-produced mucins to Siglec-9. Biochem. Biophys. Res. Commun. 402, 663–669 (2010).

55. Brinkman-Van der Linden, E. C. M. & Varki, A. New Aspects of Siglec Binding Specificities, Including the Significance of Fucosylation and of the Sialyl-Tn Epitope. J. Biol. Chem. 275, 8625–8632 (2000).

56. Paulson, J. C., Macauley, M. S. & Kawasaki, N. Siglecs as sensors of self in innate and adaptive immune responses. Ann. N. Y. Acad. Sci. 1253, 37–48 (2012).

