## Supplemental Information for "Immune-informed mucin hydrogels evade fibrotic foreign body response *in vivo*"

### Supporting information

### Materials and methods

#### Materials

Amine-derivatives of tetrazine (Tz) and norbornene (Nb) were purchased from Bioconjugate Technology Company and TCI Europe N.V., respectively. All other chemicals were purchased from Sigma Aldrich. PCR related reagents were purchased from ThermoFisher Scientific. RNA extraction micro or mini kits were purchased from Qiagen.

#### Synthesis of BSM Tz and Nb

Bovine submaxillary mucin (BSM) was pre-dissolved in MES buffer (0.1 M MES, 0.3 M NaCl, and pH 6.5) at the concentration of 10 mg/mL. To this reaction mixture, 1-ethyl-3-(3-dimethylaminopropyl) carbodiimide (EDC; 4 mmol per gram of dry mucin) and Nhydroxysuccinimide (NHS; 4 mmol per gram of dry mucin) were added and stirred for 15 min. Thereafter, 1 mmol of tetrazine-amine (Tz) and 2 mmol of norbornene-amine (Nb) were added individually to generate the BSM tetrazine (BSM-Tz) or BSM norbornene (BSM-Nb). The reaction mixtures were stirred overnight at 4 °C and then dialyzed in MWCO (10 kD cutoff) dialysis tubing for 2 days against 300 mM NaCl and then MilliQ (MQ) $H_2O$ for 2 day. The samples were freeze-dried and stored in -20°C.

**NMR analysis of the Tz and Nb functionalities**

BSM derivatives were dissolved in deuterium oxide (Sigma-Aldrich). After solubilization, 600 μL of the solutions were transferred into a 5 mm NMR tube (Norrel, USA). $^1$H-NMR spectra were obtained on a Bruker Ultrashield plus 500 MHz spectrometer (Bruker Corporation, USA). The processing for the spectra was carried out by using the MestReNova software (version 12.0.1-20560).

**Examining the sialic acid (Neu5Ac) content in BSM and BSM derivatives.**

The conjugation of Tz and Nb used in this study were achieved by targeting the activated carboxylic groups on BSM. To locate whether the functionalities are on the BSM protein backbone or the tips of O-glycans sialic acid. We examined the sialic acid content in BSM and BSM-Tz and BSM-Nb via high-performance anion-exchange chromatography (HP-AEC) based method. In brief, we solubilized the BSM, BSM-Tz, and BSM-Nb (2.5 mg in 500 μL MQ $H_2O$) and then added 500 μL of sulphuric acid (0.1 N) and incubated at 80 °C for 1 hour to release the sialic acid. This method has been proven can remove maximum sialic acid content from BSM[1]. 500 μL of sodium hydroxide (0.1 N) was then added to the solutions to neutralize the pH. Neu5Ac (Sigma Aldrich) dissolved in MQ $H_2O$ was used as the standard. The samples were filtered and injected (10 μl) into the CarboPac PA1 column (4 × 250 mm,

Dionex). Neu5Ac was eluted by a solution containing 33% of 300 mM NaOH and 15 % of 1 M sodium acetate in $H_2O$ and detected by pulsed amperometric detection (HPAEC-PAD) with an ICS-3000 system (Dionex)[2]. The Neu5Ac content was calculated via the integration of sialic acid elution peak in comparison to sialic acid standard curve.

**Rheological characterization of BSM hydrogels**

Rheological measurements were performed using a commercial shear rheometer (MCR302, Anton Paar) equipped with a plate-plate measuring geometry (measuring head: PP25, Anton Paar, Graz, Austria). The gap between the measuring head and the bottom plate (P- PTD200/Air, Anton Paar) was set to $d$ = 150 µm for all measurements. Immediately before measurement, the two components of the Muc-gel (BSM-Tz and BMS-Nb) were diluted in the PBS (pH 7.4) to a concentration of 25 mg/mL each. The two components were thoroughly mixed and centrifuged to remove bubbles before 100 µL of the sample were pipetted onto the rheometer plate. First, gel formation was analyzed for a total time span of $t$ = 100 min. Both, the storage ($G'$) and loss modulus ($G''$) were determined by a torque controlled ($M$ = 5 µNm) oscillatory ($f$ = 1 Hz) measurement. Afterwards, a strain-controlled frequency sweep (from $f_{\text{start}}$ = 10 Hz to $f_{\text{end}}$ = 0.01 Hz) was performed to determine the frequency dependent viscoelasticity of the cross-linked sample. For this frequency sweep, a constant strain was used which was chosen as the average of the five last values determined from the prior torque-controlled measurement.

To test the stability of the mucin gel towards enzymatic degradation, we introduced a custom-made rheological setup, which allowed for exposing the sample to an enzyme solution *in situ*. This setup consists of a commercial PP25 measuring head (Anton Paar) and an in-house developed bottom plate – from now on referred to as holey plate – that can be mounted onto the bottom plate of the rheometer (P-PTD200/80-I, Anton Paar). This holey plate comprises

19 regularly orientated holes with a diameter of 1.5 mm each. This design was chosen to allow fluid to diffuse from a reservoir chamber located below into the actual sample located above the holey plate. The sample and the holey plate were separated by a polycarbonate membrane (Whatman Nuclepore 50 nm, Sigma Aldrich) to prevent the sample from leaking into the fluid chamber while allowing small molecules (such as trypsin) to penetrate the membrane and to enter the sample. An inlet and outlet to the fluid chamber allowed for an continuous renewal of fluid during the measurement (**Scheme**), e.g., to maintain a constant concentration of enzyme in the fluid chamber.

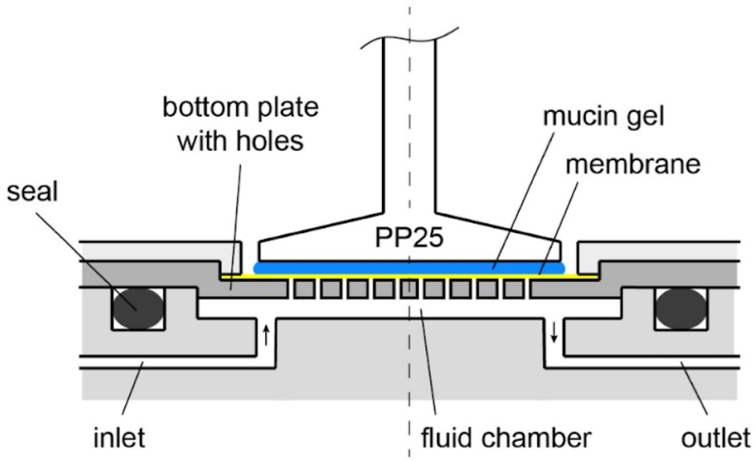


**Scheme**: Holey-Plate measuring configuration. The measuring setup consist of a commercial PP25 measuring head and a custom-made bottom plate. This bottom plate, on the one hand, allows for a fluid to diffuse from a reservoir chamber through a membrane into the sample and, on the other hand, prevents the sample from leaking into the fluid chamber.

The sample was prepared as described above; however, the measuring gap was adjusted to $d = 300$ µm to ensure that the measuring head did not interfere with the membrane. Before each measurement, the membrane was wetted and the reservoir was filled with PBS to avoid air bubbles in the system. The inlet was connected to a syringe, which was filled with PBS and then fixed to a syringe pump (LA 100, Landgraf Laborsysteme HLL GmbH, Langenhagen,

Germany). Afterwards, 200 µL of the sample was placed onto the membrane, and the measuring head was lowered to the measuring position. The measurement was performed in torque-controlled mode ($M$ = 10 µNm) and started after a delay of $t_{delay}$ = 30 min to allow the sample to cross-link. After the storage modulus had reached a steady state, the syringe pump was set to a continuous flow rate of $Q$ = 25 µL/min to equilibrate the system. After another 30 min, the buffer syringe was exchanged by a syringe filled with a trypsin solution (100 µg/mL trypsin in PBS).

**Hydrogel preparation for implantation**

All the reagents for the hydrogels preparation were sterile-filtered using a 0.45 µm pore filters. Hydrogels were prepared inside a cell culture laminar hood. Alginate hydrogels (Alg-gel) were formed by mixing 2.5 % solution of clinical grade alginate (PRONOVA SLG20, NovaMatrix) dissolved in 0.9 % saline solution (pH 7.4), and crosslinked with $BaCl_2$ gelling solution (20 mM $BaCl_2$, 250 mM D-mannitol, 25 mM HEPES, n ($Ca^{2+}$) = n (monomer of guluronate)). BSM hydrogels were prepared by mixing equal volume of 2.5 % BSM-Tz and BSM-Nb dissolved in PBS. The hydrogel were allowed to reach gelling equilibrium then immersed in a saline solution. For each hydrogel, 50 µL disks were prepared using a 1 mL syringe with the tip cut off (BD Bioscience). After formation, the alginate disks were washed with HEPES buffer (25 mM HEPES, 1.2 mM $MgCl_2\times 6H_2O$, 4.7 mM KCl, 132 mM $NaCl_2$). Alg-gels were then kept at 4 °C before the implantation in a saline solution, while the BSM disks were stored in saline solution. All the solutions used for alginate hydrogel formation have a pH of 7.4 and an osmotic pressure of 290 mOsm. 2 independent experiments were performed in 4 or 5 different mice. In each mouse, 4 gels were implanted.

**Biocompatibility assay**

The biocompatibility of Muc-gel was investigated by alamar blue assay. In brief, monocytes cell line THP-1 was maintained in a complete RPMI 1640 culture medium containing 10 % FBS, and 1 % penicillin/streptomycin and incubated in a humidified incubator with 5 % $CO_2$, at 37$^{o}$C. THP-1 cells were then incubated with the complete culture medium supplemented with 150 nM phorbol-12-myristate-13-acetate (PMA, Sigma, P8139-Avoid light) for 3 days followed by 24 h incubation in the complete cell culture medium to obtain THP1-derived macrophage type 0 (M0). To embed M0 cells in 3D Muc-gel, cells were resuspended in 2.5 % BSM-T/N solution which was pre-solubilized in the complete culture medium. The Muc-gel cylinder was then prepared as described above. M0 cells in 3D Muc-gels were cultured for 7 days in a humidified incubator with 5 % $CO_2$, at 37 $^{o}$C. Alamar blue assay was performed on day 0, 1, 3, and 7. Cells were incubated with 10 % resazurin in the complete culture medium for 4 h, and the fluorescent intensity was measured by CLARIOStar microplate reader at the excitation of 530 nm and emission of 590 nm. The fluorescence intensities measured on day 1, 3, and 7 were normalized to the fluorescence intensity measured on day 0. Three independent experiments were performed.

**Hydrogel implantation**

Mice were housed under standardized conditions (21-22°C, 12-h light and 12-h dark cycle) in SPF facility and were used at 10-12 weeks old under guidelines of the Swedish National Board for Laboratory approved by the Swedish Laboratory Animal Ethical Committee in Uppsala.. Mice were allowed to acclimatize for one week prior to the surgery. For peritoneal cavity (i.p.) implantation of the hydrogels, the mice were anesthetized by 2.5% of isoflurane and the fur on the abdomen was shaved and sprayed with 70% ethanol. 1 cm length incision was done to implant the hydrogels in peritoneal cavity and then sutured.

**Isolation of peritoneal and gel-infiltrated cells**

After 14 and 21 days, the mice were euthanized by isoflurane followed by cervical dislocation. Cells were obtained by lavage from the peritoneal cavity and by isolation from the harvested hydrogels. 5 mL of ice-cold PEB solution (1X PBS, pH 7.4, 2 mM EDTA, and 0.5% BSA) were injected into the peritoneal cavity in order to collect the free-floating cells. The implanted hydrogels were harvested and placed into a 5 mL PBS solution and single-cell suspension was prepared using a gentleMACS Dissociator (Miltenyi Biotec) following by the manufacturer's instructions. Cells were then filtered through 70 μm cell strainer (Fisher Scientific) and the red blood cells were lysed by hypotonic shock (0.2% NaCl, 25 s, v/v 1.6% NaCl).

**Flow cytometry cell sorting**

Cells were incubated with mouse BD Fc-block (BD biosciences, 2.5 μg per $1x10^6$ cells in 100 μL) at 4$^o$C for 15 min to avoid nonspecific antibody binding and then stained with antibody cocktail in dark at 4$^o$C for 30 min. The antibody cocktail contains the following fluorescent conjugated monoclonal antibodies: CD68 (1 μL per 1 million cells in 100 μL staining volume, CD68-Alexa647, Clone: FA-11, Cat. No. 137004, BioLegend); CD11b (1.25 μL per 1 million cells in 100 μL staining volume, CD11b-PE, Clone: M1/70, Cat. No. 101207, BioLegend); Gr-1 (0.5 μL per 1 million cells in 100 μL staining volume, CD11b-Alexa488, Clone: RB6-8C5, Cat. No. 108417, BioLegend); CD19 (1.25 μL per 1 million cells in 100 μL staining volume, CD19-PE/Cy7, Clone: 6D5, Cat. No. 115520, BioLegend); IgM (5 μL per 1 million cells in 100 μL staining volume, IgM-BV421, Clone: RMM-1, Cat. No. 406532, BioLegend). Cells were washed with 3 mL PBS twice before to be resuspended in the PBS with 1% BSA and to be sorted (BD FACSAria™, BD Bioscience). Data were analysed with Flowjo 10.5.2 software (Tree Star, Inc).

**Gene expression analysis by real-time PCR**

Total RNA of cells was extracted by using either Qiagen RNeasy mini kit or Qiagen RNeasy micro kit depending on cell numbers sorted. The extracted mRNA was diluted to a concentration of 0.67 ng/μL and synthesized into cDNA using Superscript III polymerase (Invitrogen). Real-time PCR was then performed to analyze the gene expression by using a TaqMan Gene Expression Master Mix (Thermo Fisher Scientific) together with TaqMan probes. The RT-PCR was carried out in a CFX96 Touch™ Real-Time PCR Detection System (Bio-Rad) with the following cycling conditions: 50°C for 2 min, 95°C for 10 min, 95°C for 15 sec, 60°C for 1 min, and then go to step 3 for 50 cycles. Target gene relative expression to housekeeping gene Actβ were performed. The validation of housekeeping gene was performed for both RL37 and Actβ and revealed that Actβ showed better consistency among tested samples.

**Histological staining**

Muc-gel and Alg-gel explants were fixed by 4% paraformaldehyde overnight and then washed by 70 % ethanol, before being processed for paraffin embedding. Samples were cut into 5 μm section, and then deparaffinized, and rehydrated for Hematoxylin and Eosin (H&E) and Masson's trichrome staining. The slides were then dehydrated and mounted with cover slides. The images were scanned by using automated image scanner (Aperio scanscope AT, Leica Biosystems Imaging, Inc.) and analyzed by Aperio ImageScope software (Leica Biosystems Imaging, Inc.).

**Immunohistofluorescent study**

Immunofluorescent imaging was performed to examine the expression of TGFβ-1 and α-smooth muscle actin. In brief, deparaffinized and rehydrated paraffin sections of Muc-gels and Alg-gels were firstly washed with PBS buffer. Sections were permeabilized for 30 mins with 0.1 % Triton X-100 solution followed by 3 times wash with PBS. Sections were then blocked by incubation with 1% bovine serum albumin (BSA) solution for 1 h followed by 3 times wash with PBS buffer. Sections were incubated with primary anti-TGFβ-1 antibody (Novus Biologicals, Cat. No. AB-246-NA, 5 μg/mL), and anti-α-smooth muscle actin-Cy3 conjugated antibody (Sigma Aldrich, Cat. No. C6198, 1:200) overnight at 4°C and Primary anti-TGFβ-1 antibody was then labeled with secondary antibody (Alexa Fluor 488, Jackson ImmunoResearch, Cat. No. 705-485-147, 1:400). Specimen were then mounted with cover slides with mounting medium Fluoroshied™ containing DAPI (Sigma Aldrich). Images were acquired with inverted Nikon Eclipse Ti fluorescence microscope (Nikon Instruments Inc.).

**Statistical analysis**

Data are shown as mean of implants from four mice per time point and per experimental group. The significance was analysed via One-way ANOVA with Bonferroni-correction using GraphPad Prism 7.0; '*', '**', '***', and '****' indicates p value < 0.05, 0.01, 0.0005, and 0.0001. respectively.

Supplementary results

### NMR examining the functionalities of Tz and Nb

Aromatic protons in BSM-Tz spectra (δ 10.4, δ 6.0, and δ 5.2) and alkene protons in BSM-Nb spectra (δ 6.3-5.8) were detected (**Figure S-1**).

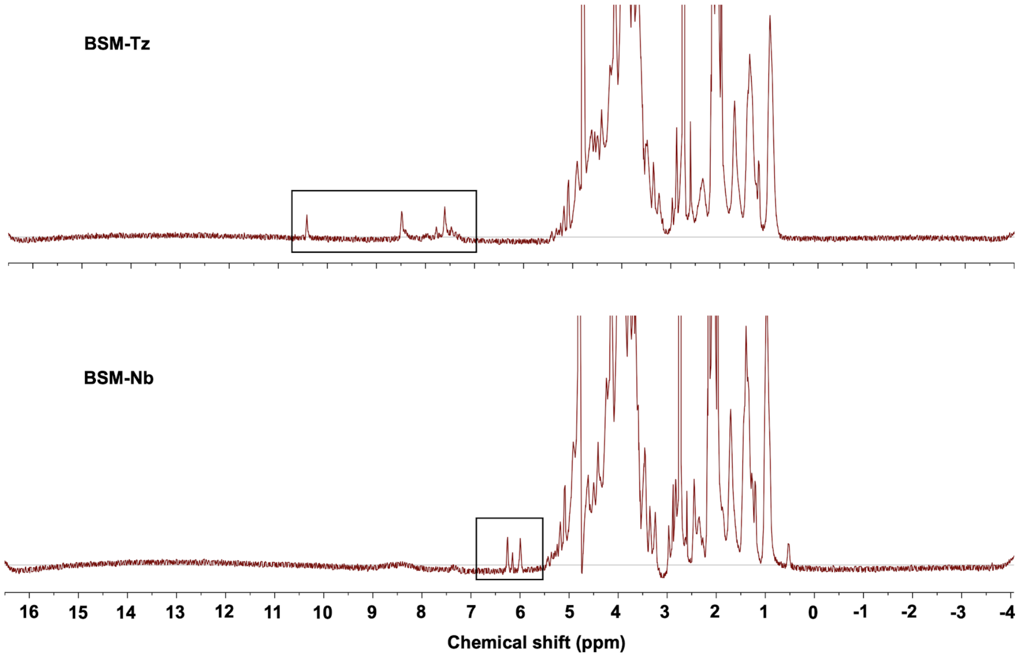


**Figure S-1.** $^{1}$H-NMR spectra of Tz and Nb functionalities on BSM. Black box indicates the appearance of aromatic protons in BSM-Tz spectra and alkene protons in BSM-Nb individually.

### Rheological property of the Muc-gel

The high shear stiffness reached here (~ 10 kPa) is already a good indication that the system is efficiently cross-linked as entangled mucin solutions only reach *G'* values on the order of 1-10 Pa[3]. The shoulder we observe (**Figure 1C, insert, dotted black line square**) in the loss modulus at *t* = 5- 9 min could indicate the existence of two different reaction mechanisms, which occur sequentially during the cross-linking process. Even though the system reaches an elastically dominated state, the contribution of viscous effects is still significant as indicated by the relatively high *G''* values. This is typical for networks comprising (semi-)flexible polymers and indicates the presence of some form of dissipation mechanism e.g., from thermal polymer fluctuations in a viscous medium (water/buffer).

### Calculation of the Muc-gel pore size

We further estimated the average molecular weight (□□ , the molecular weight of chain segments between two adjacent crosslinks or entanglement points ) is ~ 6,17 kg/mol. Since the hydrogels showed elastic character, we applied rubber elasticity theory to approximate the mesh size (□, the distance between two adjacent crosslinks or entanglement points, ~ 7,4 nm) and revealed our hydrogels would allow the non-restricted transportation for nutrients, metabolism products, and secretones by the cells[4]. The following equations[5–7] were used:

$$Mc = \frac{c\rho RT}{G'_P}$$

$$□ = \left(\frac{□'_{□}□_{□}}{□□}\right)^{-1/3}$$

where □ is the concentration of polymers (2.5% w/v), $\rho$ is the density of water at 298 K (997 kg.m-3), □ is molar gas constant ($8.3144598\times10^{6}$ $cm^{3}PaK^{-1}mol^{-1}$) , $□'_{□}$ is the peak value of elastic modulus, $□_{□}$is the Avogadro constant and T is temperature (298 K).

### Cell viability study

To investigate the biocompatibility of Muc-gels, we embedded M0 macrophages derived from human monocyte cell line THP-1 in three dimensional Muc-gels and cultured them over a period of 7 days. We evaluated cell viability by alamar blue assay, in which the living cells reduce resazurin (blue, oxidized form) into resorufin which is red and highly fluorescent. As shown **Figure S-2**, the Muc-gels showed no sign of cytotoxicity over this timeframe. .

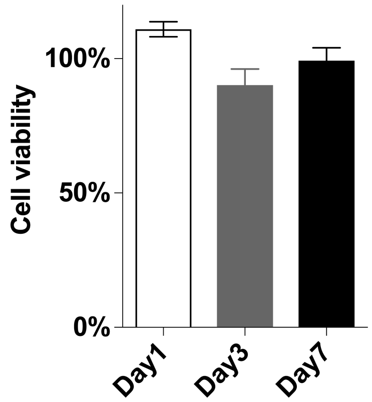


**Figure S-2.** Cell viability of THP-1-derived M0 macrophages cultured inside Muc-gel was assessed by alamar blue assay (n=3).

**Sialic acid content analysis**

The sialic acid content on BSM, BSM-Tz, and BSM-Nb eluted at the same time as control (**Figure S-3A**, **B**) and resulted in similar peak integral (**Figure S-3C**), suggesting the functionalities of Tz and Nb are not localized on the sialic acid. We rather hypothesized that they have reacted with activated carboxylic group from amino acid side chains on the mucin protein backbone.

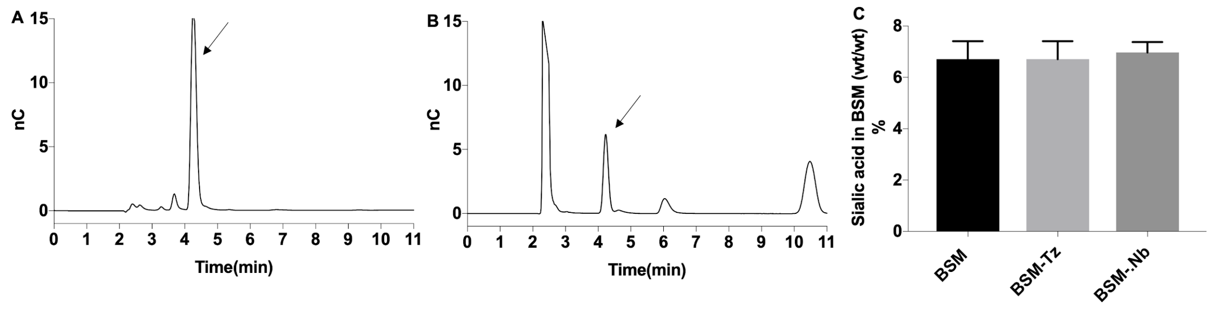


**Figure S-3.** Sialic acid content on BSM, BSM-Tz, and BSM-Nb were the same as examined by anion-exchange chromatography. The representatives of elution peaks of sialic acid standard (A), released sialic acid (B), and sialic contents in BSM and derivatives (C).

**Gene expression in myeloid (CD68$^-$Gr1$^{low/-}$CD11b$^+$) cells**

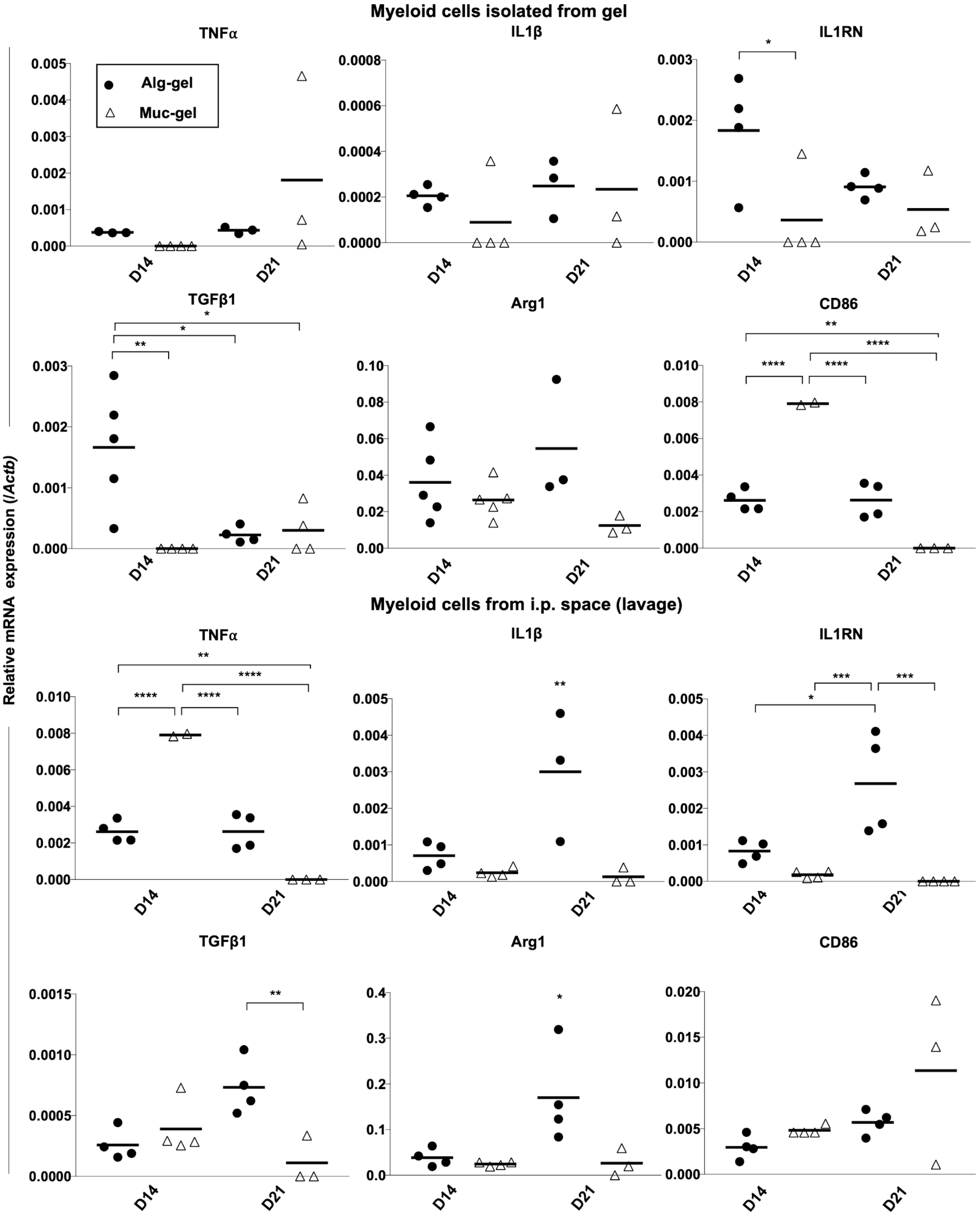


**Figure S-4.** Gene expression in sorted immature myeloid cells (CD68$^-$Gr1$^{low/-}$CD11b$^+$) from explanted Muc-gels and Alg-gels at days 14 and 21 was analyzed by RT-PCR. Data represents the mean of duplicates of RT-PCR. Data represents the mean of duplicates of RT-

PCR from 3 to 5 independent experiments and from four mice. Differences were determined using a One-way ANOVA with Bonferroni-correction.

**Reference:**


1. Varki, A. & Diaz, S. The release and purification of sialic acids from glycoconjugates: Methods to minimize the loss and migration of O-acetyl groups. *Anal. Biochem.* **137**, 236–247 (1984).
2. McKee, L. S. *et al.* A GH115 α-glucuronidase from Schizophyllum commune contributes to the synergistic enzymatic deconstruction of softwood glucuronoarabinoxylan. *Biotechnol. Biofuels* **9**, 2 (2016).
3. Biegler, M., Delius, J., Käsdorf, B. T., Hofmann, T. & Lieleg, O. Cationic astringents alter the tribological and rheological properties of human saliva and salivary mucin solutions. *Biotribology* **6**, 12–20 (2016).
4. Cruise, G. M., Scharp, D. S. & Hubbell, J. A. Characterization of permeability and network structure of interfacially photopolymerized poly(ethylene glycol) diacrylate hydrogels. *Biomaterials* **19**, 1287–1294 (1998).
5. Eiselt, P., Lee, K. Y. & Mooney, D. J. Rigidity of two-component hydrogels prepared from alginate and poly (ethylene glycol)- diamines. *Macromolecules* **32**, 5561–5566 (1999).
6. Welzel, P. B. *et al.* Modulating Biofunctional starPEG Heparin Hydrogels by Varying Size and Ratio of the Constituents. *Polymers* **3**, 602–620 (2011).
7. Yan, H. J. *et al.* Synthetic design of growth factor sequestering extracellular matrix mimetic hydrogel for promoting in vivo bone formation. *Biomaterials* **161**, 190–202 (2018).
